# RiboTRIBE: Monitoring Translation with ADAR-meditated RNA Editing

**DOI:** 10.1101/2021.06.20.449184

**Authors:** Weijin Xu, Katharine Abruzzi, Michael Rosbash

**Affiliations:** HHMI/Department of Biology, Brandeis University, Waltham, MA 02453, USA

## Abstract

RNA translation is tightly regulated to ensure proper protein expression in cells and tissues. Translation is often assayed with biochemical assays such as ribosome profiling and TRAP, which are effective in many contexts. These assays are however not ideal with limiting amounts of biological material when it can be difficult or even impossible to make an extract with sufficient signal or sufficient signal:noise. Because of our interest in translational regulation within the few *Drosophila* adult circadian neurons, we fused the ADAR catalytic domain (ADARcd) to several small subunit ribosomal proteins and assayed mRNA editing in *Drosophila* S2 cells. The strategy is named RiboTRIBE and is analogous to a recently published APOBEC-based method. The list of RiboTRIBE-edited transcripts overlaps well with ribosome profiling targets, especially with more highly ranked targets. There is also an enriched number of editing sites in ribosome-associated mRNA comparing to total mRNA, indicating that editing occurs preferentially on polyribosome-associated transcripts. The use of cycloheximide to freeze translating ribosomes causes a substantial increase in the number of RiboTRIBE targets, which is decreased by pretreating cells with the chain terminating drug puromycin. **NOTE: Additional experiments performed after first submitting this manuscript to BioRxiv estimate that less than 5% of Rps28b-ADAR is ribosome-associated. This is because the vast majority of the fusion protein sediments at the top of a polyribosome gradient. We therefore suggest that most editing reported in the manuscript is not catalyzed by ribosome-associated ADAR (10/2/2021).**

## Introduction

Cells maintain protein levels by controlling the ratio between gene expression and protein degradation. Gene expression is a multi-step process, which is tightly regulated at many levels (Gerstberger et al., 2014; Jansen, 2001; Szostak and Gebauer, 2013; Witten and Ule, 2011; Zhao et al., 1999). Translation is a crucial step, and its proper regulation is an essential and powerful way to change this equilibrium (Hershey et al., 2012). Two major factors contribute to protein synthesis rate, the abundance of an mRNA species and its translational efficiency. Translational efficiency reflects how many copies of a protein can be made from one mRNA molecule per unit time, and mRNA abundance measures how many copies of the mRNA are available in the cell to be translated.

Measuring mRNA abundance is a mundane task in modern molecular biology, as there are dozens of techniques available for this purpose, e.g., qRT-PCR, RNA-seq and state-of-the-art single cell sequencing. However, measuring translational efficiency is more complicated.

Translational efficiency is often estimated by biochemical methods like ribosome profiling (ribosome footprinting, Ingolia et al., 2009; Ingolia et al., 2011) and <u>T</u>ranslating <u>R</u>ibosome <u>A</u>ffinity <u>P</u>urification (TRAP, Heiman et al., 2014; Heiman et al., 2008; Mellen et al., 2012). Ribosome profiling is based on the fact that each ribosome covers ~28 nt of mRNA, which protects the mRNA fragment from degradation by RNase treatment (Ingolia et al., 2009). Therefore, one can estimate the translation rate of each mRNA by isolating and sequencing the mRNA fragments protected by ribosomes,. This is often done in conjunction with a separate RNA-seq experiment to normalize for mRNA abundance.

TRAP was developed as a technique to assay cell-specific translation, especially in neurons (Heiman et al., 2008). By expressing EGFP-tagged ribosome large subunit protein L10a in specific cells, TRAP uses antibody based purification of tagged ribosomes to identify mRNAs that are undergoing active translation (Heiman et al., 2014).

Although both methods have been successfully used in various contexts (Bertin et al., 2015; Iwasaki et al., 2016; Iwasaki and Ingolia, 2017; Ng et al., 2016), these biochemical assays are not ideal with limiting amounts of biological material. This is because it can be difficult to make cell extracts efficiently, perform biochemical treatments (ribosome recovery or antibody precipitation) and then generate sufficient signal or sufficient signal:noise ratio from small amounts of material, for example the 150 circadian neurons in the *Drosophila* brain (Helfrich-Forster, 2003).

Because of our long-standing interest in these neurons, we have been seeking a method to assay translation within them. Our recently developed TRIBE method provided us with a possible strategy to develop such a technique (McMahon et al., 2016; Xu et al., 2018). TRIBE fuses an RBP of interest to the ADARcd, which performs adenosine-to-inosine editing on RBP-bound RNA targets (McMahon et al., 2016). To extend TRIBE to the ribosome and translational regulation, we reasoned that fusion of the ADARcd to a ribosome protein instead of an RBP might edit actively translating mRNAs. Moreover, heavily translated mRNAs will more frequently encounter the ADARcd, which should lead to a higher chance of editing or a higher editing percentage for this mRNA population. We named this method RiboTRIBE. The fusion protein can be co-expressed in specific neurons along with a fluorescent protein, allowing FACS-based purification of the target neurons (Abruzzi et al., 2015; Rahman et al., 2018). With one RNA-seq experiment of collected cells/neurons, we can not only determine mRNA abundance but also identify editing events on different mRNAs as a measurement of translational efficiency. The strategy is identical to the recently published Ribo-STAMP method except for its use of the editing enzyme APOBEC (Brannan et al., 2021).

To search for an optimal ribosomal protein, we made constructs of several ribosome small subunit proteins fused to the ADARcd and expressed them individually in *Drosophila* S2 cells. Small subunit proteins were chosen because of their proximity to the translating mRNAs, comparing to the large subunit proteins currently in use for TRAP; they should be further from the nascent peptide chains. Our current data indicate that RPS28-ADARcd edits mRNA sufficiently to generate a credible list of translating mRNAs (Fig. 1). Indeed, several pieces of evidence indicate that RiboTRIBE edits ribosome-associated mRNAs rather than free mRNAs in the cytosol (Fig. 2, 3).

**Figure 1.**
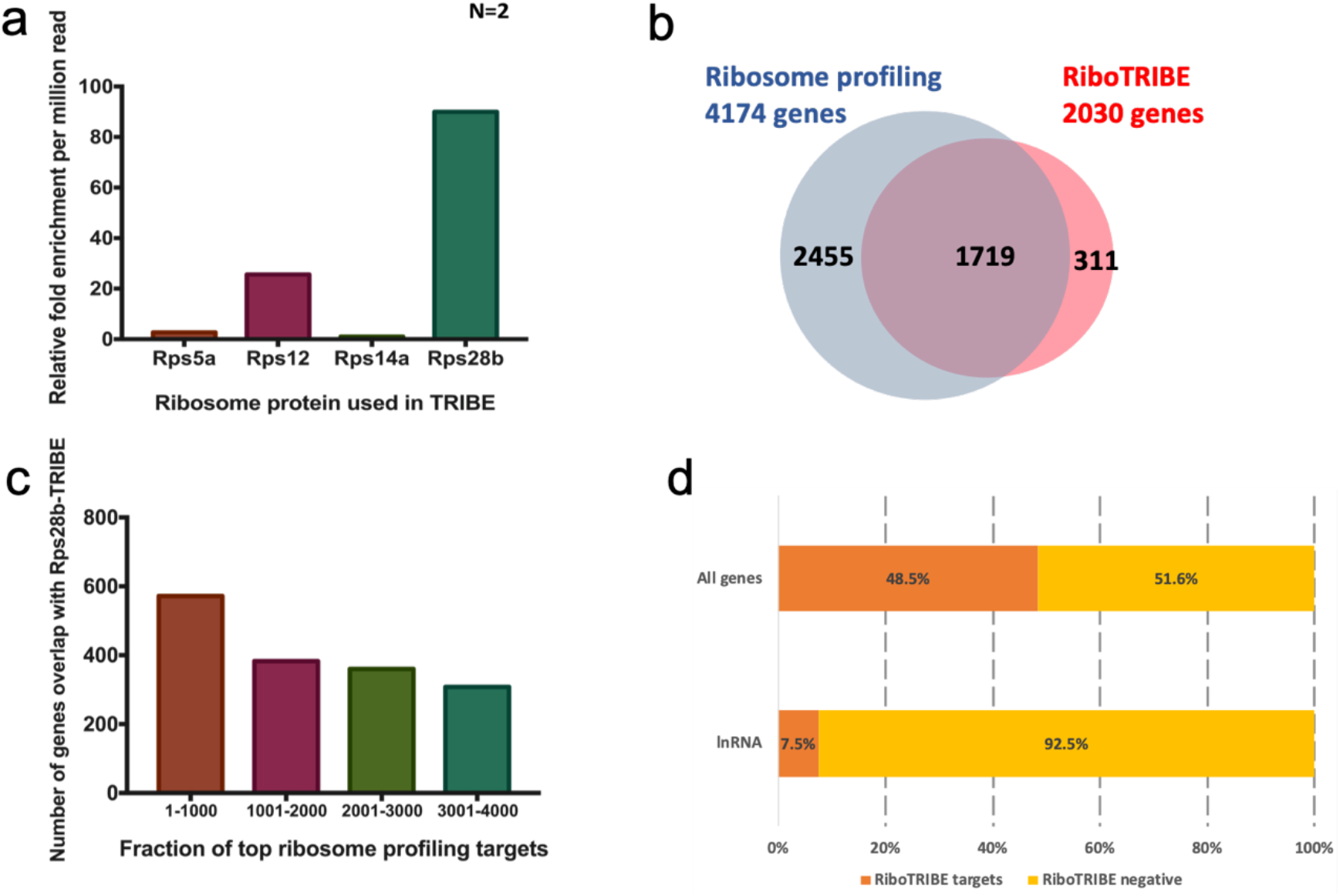
RiboTRIBE identifies a list of translating mRNAs, consistent with ribosome profiling data. (a) Both Rps12-ADADcd and Rps28b-ADARcd induced a large number of editing sites in S2 cells, but there are more (3-4 fold) identified for Rps28b. Rps28b-ADARcd was used for all further experiments. Background editing sites in wild-type S2 cells were subtracted from the output. The number of sites identified are normalized to the sequencing depth of each sample and are measured by relative fold change compared to the lowest sample Rps14a-ADARcd (see also Chapter 2, method). (b) The genes identified by RiboTRIBE were compared to those genes identified via ribosome profiling from the same S2 cells. 85% of the 2030 RiboTRIBE identified genes were also identified in ribosome profiling studies (Z-test performed, p<0.00001). (c) Histogram shows how many of the top ranked ribosome profiling targets were also identified using RiboTRIBE genes. 60% of top 1000 ribosome profiling targets (brown) are reproduced by RiboTRIBE, and the percentage gradually decreases for the lower ranked ribosome profiling targets. (d) A comparison of the number of highly expressed genes or lncRNAs that were identified as RiboTRIBE targets. Nearly 50% of the highly expressed genes were identified by RiboTRIBE. In contrast, only ~7% of lncRNAs were edited by RiboTRIBE. This difference is statistically significant (Z-test performed, p<0.00001).

**Figure 2.**
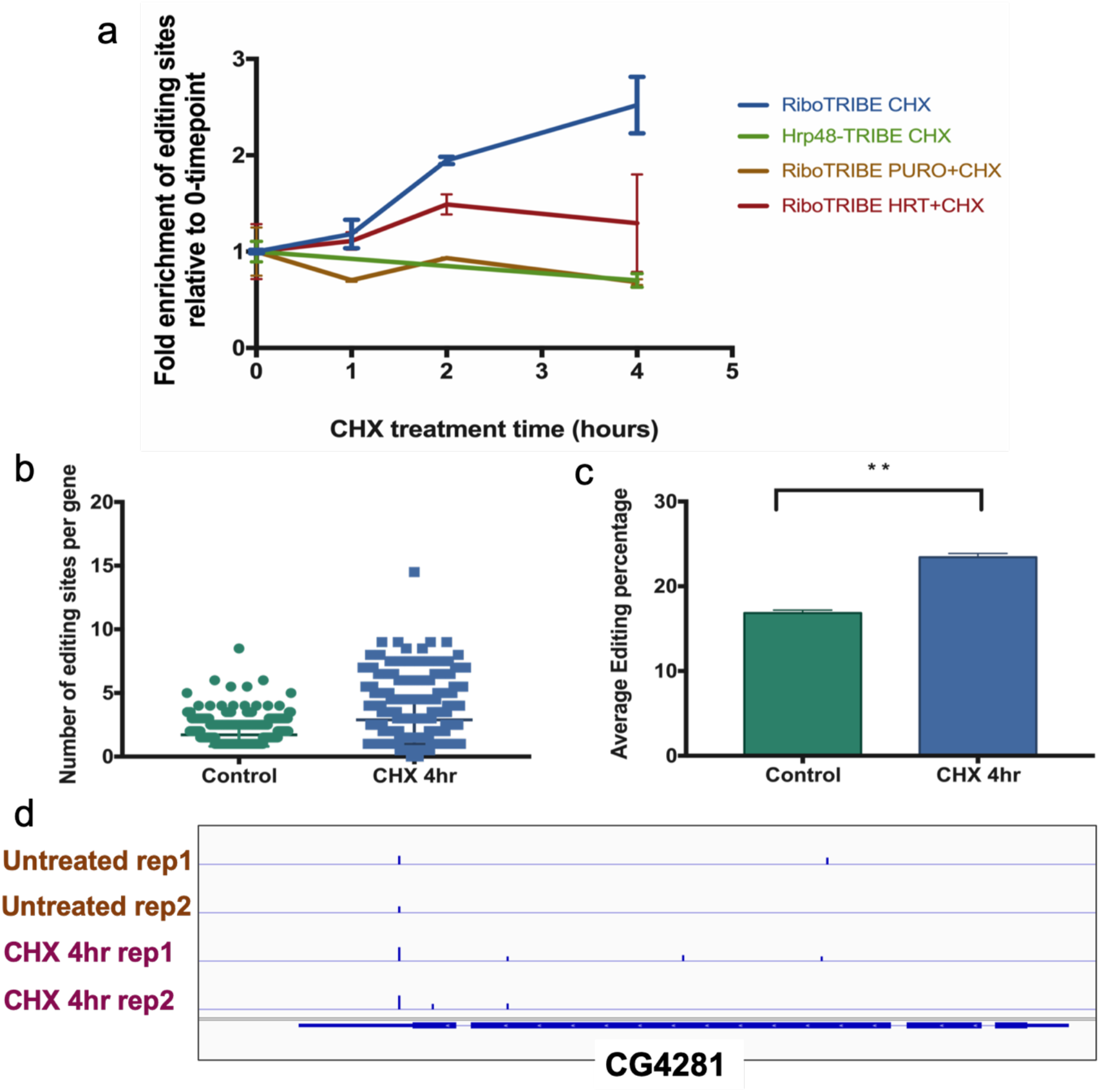
Cycloheximide treatment of RiboTRIBE expressing cells induces an increase in the number of editing sites, which is negated by puromycin pretreatment. (a) Graphs showing the fold change of the number editing sites after CHX treatment of 1 hour, 2 hours and 4 hours. Cells were allowed the standard 24 hours of RiboTRIBE expression prior to the treatment. RiboTRIBE expressing cells were treated with cycloheximide (CHX, blue), 30 min of puromycin before cycloheximide (PURO+CHX, yellow) and 30 min of harringtonine before cycloheximide (HRT+CHX, red). Hrp48 TRIBE expressing cells were treated with cycloheximide serves as the negative control (green). The number of sites identified were normalized to the sequencing depth of each sample and were measured by relative fold change compared to the 0-time point data of each condition (see also Chapter 2, method). Error bar showns as the standard deviation. (b) Dot plot showing the number of editing sites per gene identified by RiboTRIBE without treatment (green) and RiboTRIBE after 4 hours of CHX incubation (blue). The average number of sites per gene was 1.7 vs 2.9, significantly higher after CHX incubation (t-test, p<0.0001). (c) The average editing percentage of the editing sites shared between RiboTRIBE increases with 4 hours of CHX incubation (blue) compared to the same cells without treatment (green).. The average editing percentage are 17% vs 23.5%, significantly higher after CHX incubation (paired t-test, p<0.0001). (d) A screen shot of the IGV genome browser shows the editing sites identified by RiboTRIBE without treatment (top two lanes) and RiboTRIBE with 4 hours of CHX incubation (bottom two lanes) on the gene, CG4281. Each blue bar is an editing site identified and its height represents the editing percentage.

**Figure 3.**
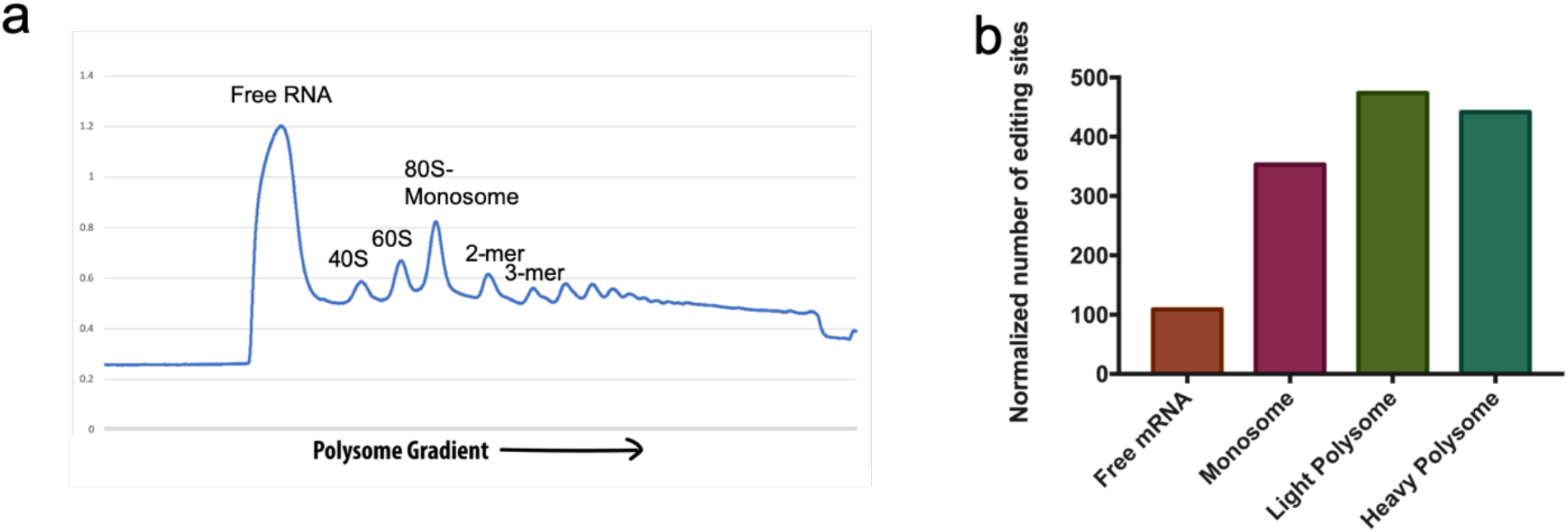
Polysome gradients performed on RiboTRIBE expressing cells reveal that RiboTRIBE editing sites are more enriched in the ribosome bound fraction of the gradient. (a) OD 264nm reading of the polysome gradient after ultracentrifuge. Free RNA fraction fraction, 80S fraction, 2 and 3 polysome (as light polysome) and higher polysome (as heavy polysome) were collected respectively. (b) Bar graph showing the number of editing sites induced by RiboTRIBE from each fraction of the polysome gradient. The number of sites identified are normalized to the sequencing depth of each sample.

## Results

The ribosomal protein L10a was fused to GFP for TRAP (Heiman et al., 2008), and our lab previously attempted to fuse this ribosomal protein with the ADARcd and express the fusion protein in S2 cells. However, there were very few editing events (similar to endogenous ADAR editing; data not shown), indicating that editing was inefficient with L10a-ADARcd. There are many possible explanations, but perhaps the most straightforward one is that a large subunit protein placed the ADARcd too far from the translating mRNAs. This hypothesis implies that this strategy might work if the ADARcd is fused to a ribosome protein physically closer to the translating mRNA.

From the published crystal structure of the eukaryotic 40S ribosome subunit (Weisser et al., 2013), we identified several small subunit proteins that have access to the mRNA tunnel. We also required these proteins to sit on the outer surface of the 40S subunit, so that fusion with a bulky ADARcd (~40 kDa) might not severely inhibit ribosome assembly. The final four candidates were *Drosophila* S5a, S12, S14a and S28b (Paralogs were chosen by their expression pattern in flies). After expressing RiboTRIBE constructs with each of the four ribosome proteins fused with ADARcd in *Drosophila* S2 cells for 3 days, we used the co-expressed GFP signal to FACS sort the positive transfected cells. The necessity of FACS sorting was due to the low transfection efficiency in S2 cells (typically <10% at this time in our hands), but it also conveniently imitates the situation in brains where the desired neurons are scarce. RNA-seq libraries were prepared from the collected cells, sequenced on an Illumina Nextseq 500 and analyzed computationally to identify editing events (Rahman et al., 2018). Of the four ribosome small subunit proteins tested, Rps28b-ADARcd induced the most consistent editing sites by a several-fold margin (Fig. 1a). We therefore only used this construct for subsequent RiboTRIBE experiments.

2030 of the ~6000 highly expressed S2 cell transcripts were edited by RiboTRIBE. These transcripts must contain at least one successful editing event, which requires at least 2% editing, a threshold of 20 reads at that position and be reproducible in both replicates (Fig. 1b). The lower required editing percentage here compared to RBP-TRIBE (typically 10%) accommodates the dynamic movement of translating ribosomes; they are less likely to edit associate transcripts as efficiently as a static RBP-TRIBE fusion protein.

Not surprisingly perhaps, only 30% of the RiboTRIBE editing sites were shared with Hrp48 TRIBE sites, indicating that the RiboTRIBE results are specific (data not shown). To further address the translation of the RiboTRIBE transcripts, we performed a S2 cell ribosome profiling experiment. With a threshold of RPF>5 and input FPKM>3, ribosome profiling identified 4174 transcripts undergoing translation in S2 cells. 1719 of these transcripts were shared with RiboTRIBE transcripts (~85% of the RiboTRIBE dataset, p<0.00001, Fig. 1B).

Interestingly, RiboTRIBE data agree even more strongly with ribosome profiling transcripts that are higher confidence or more heavily translated (Fig. 1c). They are identified by ranking the ribosome profiling targets by normalized signal-to-FPKM ratio. 572 of the top 1000 ribosome profiling targets are also identified by RiboTRIBE, whereas the percentage decreases for lower ranked ribosome profiling targets. These results also indicate that RiboTRIBE preferentially edits translating mRNAs.

To examine the overlap between the RiboTRIBE and ribosome profiling data another way, we performed a correlation test. We used the total editing percentage of each gene as the parameter for RiboTRIBE and the profiling score normalized to FPKM for ribosome profiling. There was a weak correlation between the two parameters (correlation coefficient = 0.15, and p-value =1.3e^−10^, Fig. S1). This presumably reflects differences in editing efficiency of different mRNAs unrelated to translational efficiency (See Discussion).

As an additional control, we examined whether RiboTRIBE distinguishes between coding mRNAs and long non-conding RNAs (lncRNAs). Of the 3000 most abundant RNA species in S2 cells, 48.5% are RiboTRIBE targets, suggesting that a substantial fraction of these RNAs are being translated at the time of assay (Fig. 1d). We noted that 40 of the 3000 transcripts are lncRNAs, which by definition should not be translated. Only 3 of these 40 (7.5%) were identified as RiboTRIBE targets, a striking decrease in ratio compared to regular mRNAs (Fig. 1d). lncRNAs are nuclear by definition, which should make them inaccessible to the translation machinery and therefore to RiboTRIBE editing. We do not have an explanation for the three outlier lncRNAs, but it is not unprecedented to find peptide synthesis from some lncRNAs (Ji et al., 2015). Moreover, most lncRNAs in *Drosophila* are understudied, so perhaps these 3 lncRNAs are not completely nuclear. In any case, further comment requires more experiments beyond the scope of this study.

To further validate that the RiboTRIBE editing is translation-dependent, we performed a series of assays with protein synthesis inhibitors. The rationale was that ribosome translocation speed (~20 codons per second) could limit ADAR-mediated editing, because of its rather slow deamination reaction rate (<10 reactions per minute on favorable translation substrates, Kuttan and Bass, 2012). Slowing or even freezing ribosomes might therefore allow the ADARcd more time to deaminate nearby adenosines and thereby catalyze more editing. The protein synthesis inhibitor cycloheximide (CHX) inhibits ribosome translocation and at high concentration effectively freezes ribosomes on translating mRNAs (Schneider-Poetsch et al., 2010). We indeed observed increasingly more editing from CHX-treated RiboTRIBE samples starting over a 4-hour period (Fig. 2a). On the contrary, CHX treatment did not induce more Hrp48 TRIBE-mediated editing; this fusion protein is a 3’ UTR binding RBP not directly related to translation (Fig. 2a).

Puromycin (PURO) is another protein synthesis inhibitor. It causes premature nascent peptide chain termination and removes ribosomes from translating mRNAs (Azzam and Algranati, 1973). If the increasing amount of editing sites from CHX-treated RiboTRIBE samples are indeed from frozen ADARcd-tagged ribosomes, pre-treating the cells with puromycin prior to CHX addition should remove these ribosomes and inhibit the extra editing. This is indeed what occurred (Fig. 2a). Pretreating the cells with the initiation inhibitor harringtonine (HRT) had a similar but less complete effect (Fig. 2a). Note also that CHX caused an increased number of editing sites per gene (Fig. 2b) and an increased average editing percentage (Fig. 2c). Please note that CHX treatment causes more editing sites on the representative transcript CG4281, and the editing percentage of sites already identified under control conditions was also increased (Fig. 2d).

A classic way to investigate translation and ribosome state is polysome gradient analysis. Because the number of FACS-sorted RiboTRIBE-expressing cells were insufficient for generating a polysome gradient (typically > tens of millions of cells), we ran two polysome gradients in parallel, one for wild type S2 cells and one for RiboTRIBE cells. We collected the fractions for RiboTRIBE cells based on the gradient pattern from wild type S2 cells (Fig. 3a). We were able to identify many more editing sites from the ribosome-bound fractions than from the free mRNA fractions, and more in the polysome fractions than the monosome fractions (Fig. 3b).

Taken together, this series of results indicate that RiboTRIBE editing sites come substantially if not predominantly from ribosome-bound mRNAs rather than from random mRNAs.

## Discussion

In this study, we applied the recently developed TRIBE technique to translational regulation, which we named RiboTRIBE. By expressing a fusion protein between the ADARcd and ribosomal protein RPS28b, we created ADARcd-tagged ribosomes. The simple idea is that the ADARcd leaves A-to-I editing marks on mRNAs during translation, which allows detection by RNA-seq. We showed that most editing indeed comes from ribosome-bound mRNAs, a conclusion that includes the predicted effects of protein synthesis inhibitors.

We determined in the course of this work that our previous editing percentage requirement for TRIBE is not necessary and that removing it would allow detection of less frequently edited targets and a better comparison between RiboTRIBE and ribosome profiling. Unlike RBPs which have defined binding sites and probably long dwell times on mRNAs, translating ribosomes are more dynamic and move rapidly down mRNAs, which probably gives the ADARcd only a short amount of time for local editing. However, editing should still happen by chance and occur at different locations and even on different molecules of the same mRNA species. We still have to minimize random editing events as well as sequencing errors, which is done by requiring that every editing site be identified in both replicates.

We found a correlation coefficient of 0.15 between RiboTRIBE data and ribosome profiling data. Although weak, the correlation is statistically very robust and indicates that considerable RiboTRIBE editing comes from translation. In fact, when we performed a similar correlation test between S2 cell TRAP data from the literature and these S2 cell ribosome profiling data, there was no correlation (data not shown).

We used in these comparisons total editing percentage as the signal output for RiboTRIBE against ribosome profiling score normalized to FPKM. Because editing percentage is calculated by the number of edited transcripts divided by the number of total transcripts, the percentage is already normalized for mRNA abundance. Although ribosome profiling as well as TRAP are both well-recognized tools for studying translational regulation, ribosome profiling may more faithfully reflect translational efficiency as TRAP is surprisingly highly correlated with mRNA abundance (Chen and Dickman, 2017). This mRNA abundance correlation is consistent with the reports that TRAP has a relatively poor signal:noise ratio due to high background (Dougherty, 2017). Given this literature, RiboTRIBE may be a more powerful tool than TRAP to study translational regulation in small numbers of specific cells.

Ribosome profiling still has advantages. For example, ribosome profiling has nucleotide level resolution and can reflect ribosome stalling events on specific codons. There is in contrast considerable uncertainty about detailed substrate selection by RiboTRIBE independent of translational efficiency. For example, substrate choice and editing efficiency are undoubtedly influenced by local sequence, structure and perhaps even RNA flexibility (McMahon et al., 2016; Xu et al., 2018). One application that should be independent of editing efficiency considerations is comparing the translational efficiency of the same transcript under different conditions, for example at different times of day; local sequence and structure should remain constant. However, we note that even this simple idea is uncertain; RNA structure could still change even between two times of day, due for example to temporal regulation of RNA binding proteins.

Nonetheless, RiboTRIBE also has advantages, compared for example to ribosome profiling. First, RiboTRIBE does not required large amounts of starting material, making it suitable for assaying small numbers of discrete cells. Second, ribosome profiling requires an independent RNA-seq experiment to normalize for mRNA abundance in its data, which is not the case for RiboTRIBE.

A quite similar APOBEC-based method for cell-specific translational profiling was just published (Brannan et al., 2021). Although this editing enzyme may be more efficient and not suffer from the same substrate selectivity issue that affects ADAR, TRIBE may still be a viable method for RBP target identification compared to STAMP. For example, Ribo-STAMP mediated C-to-U editing may generate substantial numbers of premature stop codons. The resulting effects, including mRNA stability changes via nonsense-mediated decay (NMD, Brogna and Wen, 2009), may limit the method, especially in transgenic animals. Indeed, STAMP has not yet been shown to work in transgenic animals like TRIBE (McMahon et al., 2016; Xu et al., 2018).

One concern is, how much editing is truly translation-dependent? Might the ADARcd sit on ribosomes and then edit other mRNAs that came in close proximity? This may be a bigger problem for the CHX assay than for the control condition if CHX provides a stable anchoring point for the ADARcd and then allow transient encounters with random mRNAs. This might even contribute to some of the increased signal observed in CHX. Perhaps a comparison of transcripts that go from no editing to detectable editing with CHX addition will be different than those transcripts that just increase their editing percentage.

Another source of random editing sites could come from free Rps28b-ADARcd protein. We expected that the fusion protein would not compete favorably with the wild-type protein for incorporation during ribosome assembly. We therefore used transient transfection and copper-induced expression, so that large amounts of newly synthesized Rps28b-ADARcd protein would drive the fusion protein into ribosomes. However, some of this large amount of protein might escape or overwhelm the rapid degradation mechanism that normally takes care of excess ribosomal proteins (An and Harper, 2019; Sung et al., 2016), leaving some unassembled Rps28b-ADARcd. The best approach to disprove/confirm this possibility is to perform a western blot of each polysome gradient fraction of RiboTRIBE samples, and compare fusion protein abundance in ribosome versus free cytosol fractions. Our attempts to perform this experiment were hindered by the inefficient transfection rate of S2 cells, making it difficult to collect sufficient material for western blotting (data not shown). It is also possible that some free Rps28b-ADARcd exists in the cytosol but does not contribute much to editing; this possibility is more difficult to address.

Nonetheless, the puromycin treatment result in Fig. 2a indicates that puromycin-mediated removal of ribosomes from mRNAs is substantial. One might expect the number of editing sites to decrease similarly after puromycin pretreatment. However, the Rps28b-ADAR protein was expressed for 24 hours prior to puromycin or CHX treatment. Therefore many editing events likely occurred during those 24 hours rather than after drug treatment. The fact that the number of editing sites remained similar compared to the 0-time point indicates that less additional editing occurs after puromycin treatment (Fig. 2a).

There is still a significant path ahead to turn RiboTRIBE into an efficient method that can be applied to cells and neurons from transgenic animals and not just to tissue culture cells. This is because preliminary experiments indicate that we cannot drive sufficient RiboTRIBE expression in adult *Drosophila* neurons to allow for robust editing detection (data not shown). We note that there is also no report of Ribo-STAMP success in transgenic animals, suggesting that new strategies may be generally required for this purpose.

## Acknowledgement

The authors would like to thank members of Rosbash lab, past and present, for helpful discussions and comments, especially Kate Abruzzi for editing the manuscript draft and Hua Jin for the ribosome profiling data. MR was supported by NIH grants R01DA037721 and R01AG052465.

## Materials and Methods

### Molecular Biology

pMT-RPS-ADARcd-E488Q-V5 plasmid was created from pMT-RBP-ADARcd-E488Q-V5 plasmid (Xu et al., 2018). RPS5a, RPS12, RPS14a and RPS28b were cloned from Drosophila cDNA by PCR and inserted into pMT-RBP-ADARcd-E488Q-V5 using Gibson Assembly^®^ (NEB). Transient expression of TRIBE constructs was performed by co-transfecting pMT TRIBE plasmids with pActin-EGFP into *Drosophila* S2 cells using PEI transfection protocol. Cells were allowed to recover for 72 hours after transfection to allow for the expression of GFP before sorting GFP positive cells with a BD FACSMelody™. Total RNA was extracted from the sorted cells with TRIzol™ LS reagent (Invitrogen). TRIBE protein expression was induced with copper sulfate 24 hours before FACS sorting.

Ribosome profiling experiments were performed as previously described (Jin et al, submitted).

For CHX treatments, cells were incubated with CHX for 1 hour, 2 hours or 4 hours prior to FACS sorting. Puromycin or harringtonine pretreatment of cells were performed 30 minutes before CHX addition.

NEXTFLEX^®^ Rapid Directional qRNA-seq library Kit (Perkin Elmer Inc.) was used to construct RNA-seq libraries from S2 cell RNA. All libraries were sequencing by an Illumina NextSeq^®^ 500 sequencing system using NextSeq^®^ High Output Kit v2 (75 cycles). Each sample were covered by ~20 million raw reads.

### Polysome Gradient Centrifugation

Sucrose gradients were prepared in Beckman polyallomer tubes (No. 331372) with an 11mL gradient. The gradient consists of 11 layers of 1mL sucrose solution with gradual decreasing concentrations from 1.5M, 1.4M… to 0.5M (1.5M on the bottom, 0.5M on the top). These sucrose solutions were buffered using 50mM Tris PH7.5, 0.25M KCl, and 50mM MgCl2 and 0.1mg/mL cycloheximide was added. After adding each sucrose solution layer with a 1mL serological pipet (with gravity mode on pipet aids) without disrupting the separating surface, the gradient tubes were sealed with parafilm on top and stored at −80degrees Celsius. Gradient tubes were thawed at 4degrees Celsius overnight before using.

S2 cells from one T75 flask were used for each polysome gradient. Cells were treated with 0.1mg/mL cycloheximide for 15 minutes (or other conditions as indicated). Lysis buffer was made with buffer solution as above plus 0.5% Triton X-100, protease inhibitors and 1ul/0.1mL Superasin (RNase inhibitor). Cells were pelleted, resuspended in 500uL of lysis buffer, vortexed, and lysed on ice for 15 minutes. The lysate was micro-centrifuged at max speed for 20 minutes at 4degrees Celsius before the supernatant was collected. Equal amounts of supernatant (measured by A260 absorbance) were added on top of each gradient and ultra-centrifuged for 3 hours 20 minutes at 32000 rpm at 4degrees Celsius with SW41 Ti rotor.

Fractions were collected with BR-188 Density Gradient Fractionation System and subjected to standard RNA extraction.

### RNA-editing Analysis

RNA sequencing data were analyzed as previously described (McMahon et al., 2016; Rodriguez et al., 2012; Xu et al., 2018), with minor modifications. RNA-seq data was mapped to the current dm6 genome. PCR duplicates were removed based on the UMIs on each end of the final DNA fragments. Background editing sites found in samples expressing Hyper-ADARcd alone were subtracted from the TRIBE identified editing sites in S2 cells. Overlapping of editing sites from two datasets was identified using “bedtools intersect” with parameters “−f 0.9 −r”. The criteria for RNA editing events were: 1) The nucleotide was covered by a minimum of 20 reads in each replicate; 2) More than 80% of genomic DNA reads at this nucleotide is A with zero G (use the reverse complement if annotated gene is on the reverse strand); 3) At least one G is observed at this site in mRNA (or C for the reverse strand).

Quantification of RNA sequencing reads distribution was performed with read_distribution.py script in RSeQC v2.3.7 (Wang et al., 2012).

---

**Figure S1.**
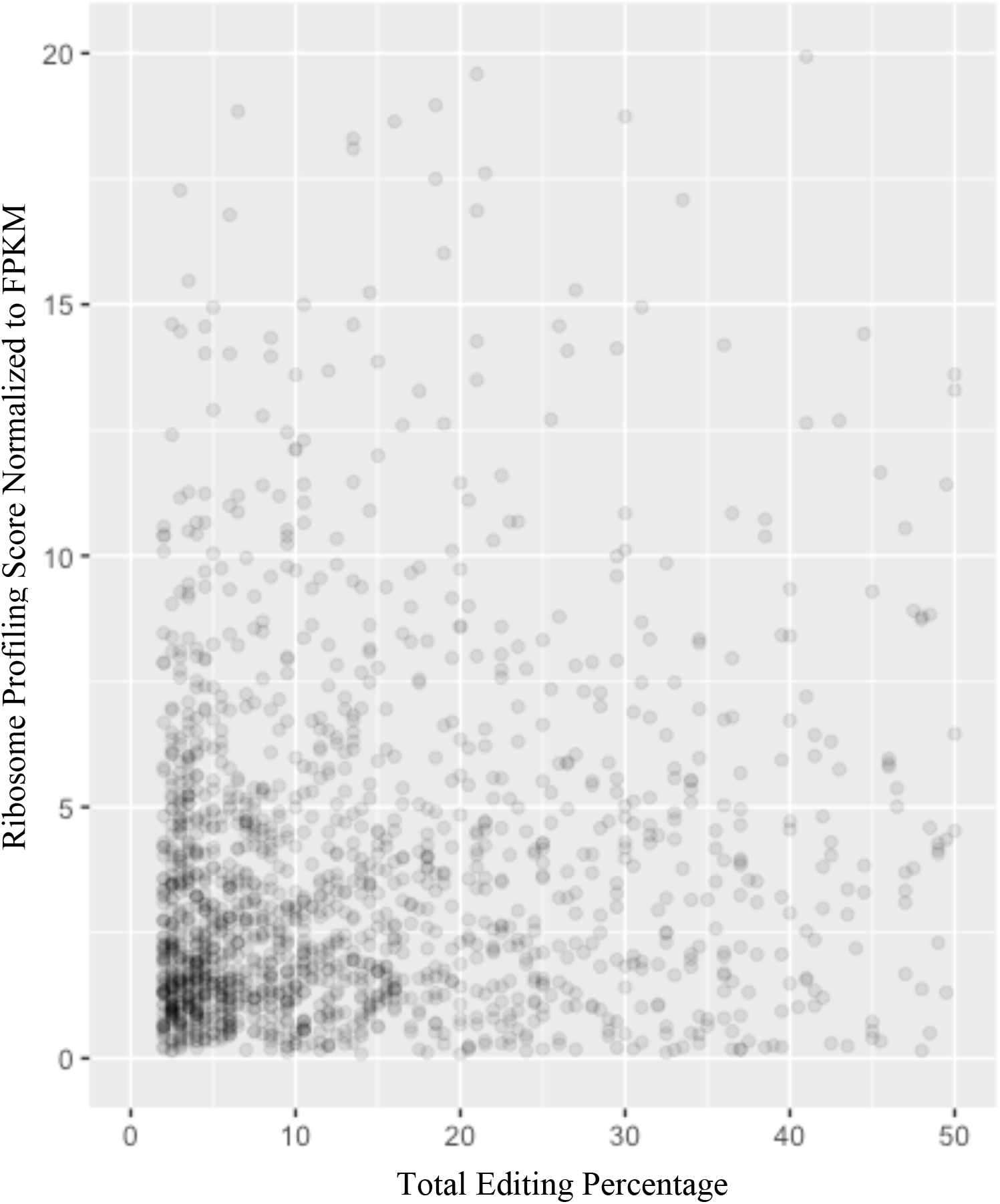
The total number of editing sites is weakly correlated with normalized ribosome profiling signal intensity. Each dot on the plot represents one gene identified by both methods. The X-axis is the total number of editing sites of each gene averaged between two replicates, and the Y-axis the RPKM signal from ribosome profiling normalized to FPKM of each gene (from an independent RNA-seq library from S2 cells). Weak correlation was observed from the two variables with the pearson method (correlation coefficient = 0.15, and p-value =1.3e^−10^).

## Notes

### Competing Interest Statement

The authors have declared no competing interest.

